# Loss of Linker Histone H1 in the Maternal Genome Influences DEMETER-Mediated Demethylation and Affects the Endosperm DNA Methylation Landscape

**DOI:** 10.1101/2022.10.17.512625

**Authors:** Qiang Han, Yu-Hung Hung, Changqing Zhang, Arthur Bartels, Matthew Rea, Hanwen Yang, Christine Park, Xiang-Qian Zhang, Robert L Fischer, Wenyan Xiao, Tzung-Fu Hsieh

## Abstract

The *Arabidopsis* DEMETER (DME) DNA glycosylase demethylates the central cell genome prior to fertilization. This epigenetic reconfiguration of the female gamete companion cell establishes gene imprinting in the endosperm and is essential for seed viability. DME demethylates small and genic-flanking transposons as well as intergenic and heterochromatin sequences, but how DME is recruited to these target loci remains unknown. H1.2 was identified as a DME-interacting protein in a yeast two-hybrid screen, and maternal genome H1 loss affects DNA methylation and expression of selected imprinted genes in the endosperm. Yet, the extent to which how H1 influences DME demethylation and gene imprinting in the *Arabidopsis* endosperm has not been investigated. Here, we showed that unlike in the vegetative cell, both canonical histone H1 variants are present in the central cell. Our endosperm methylome analysis revealed that without the maternal linker histones, DME-mediated demethylation is facilitated, particularly in the heterochromatin regions, indicating that H1-containing nucleosomes are barriers for DME demethylation. Loss of H1 in the maternal genome has a very limited effect on gene transcription or gene imprinting regulation in the endosperm; however, it variably influences euchromatin TE methylation and causes a slight hypermethylation and a reduced expression in selected imprinted genes. We conclude that loss of maternal H1 indirectly influences DME-mediated demethylation and endosperm DNA methylation landscape but does not appear to affect endosperm gene transcription and overall imprinting regulation.

## Introduction

DNA methylation regulates important processes in eukaryotic genomes including gene transcription, transposon silencing and genomic imprinting (Law and Jacobsen, 2010). In plants, *de novo* methylation is guided by small interfering RNAs in a process known as the RNA-directed DNA methylation (RdDM) (Matzke and Mosher, 2014). Once DNA methylation is established, it is maintained upon replication by distinct DNA methyltransferases to preserve cell identity and genome integrity. Plant DNA methylation is found in CG, CHG, and CHH sequence contexts (H is A, C, or T) and is mainly targeted to transposons or repetitive sequences of the genome. DNA methylation also needs to be dynamically reconfigured during development to enable transition to a new cell fate and transcriptional state. Such epigenetic reconfiguration plays a prominent role in animal reproduction and is required for reproductive success in flowering plant (Ono and Kinoshita, 2021). Removal of DNA methylation in plant is catalyzed by the DNA glycosylase DEMETER (DME), repressor of silencing1 (ROS1), DEMETER LIKE2 (DML2), and DML3 in *Arabidopsis* (Agius et al., 2006;Gehring et al., 2006) through a Base Excision Repair (BER) pathway. Whereas ROS1, DML2 and DML3 are more widely expressed and function to counteract the spread of RdDM methylation into nearby coding genes, DME demethylation during reproduction establishes gene imprinting in the endosperm and is essential for seed viability (Choi et al., 2002;Gehring et al., 2006;Gent et al., 2022;Xu et al., 2022).

During *Arabidopsis* reproduction, the central cell genome is extensively demethylated at about nine thousand loci prior to fertilization. This epigenetic reconfiguration differentiates the imprints of parental genomes and establishes the parent-of-origin specific expression of many imprinted genes, including two essential components of the PRC2 complex (i.e., *MEDEA* and *FIS2*) crucial for seed development (Choi et al., 2002;Jullien et al., 2006). In pollen, DME also demethylates the vegetative cell genome to reinforce DNA methylation of the sperm and to ensure a robust pollen germination in certain ecotypes (Schoft et al., 2011;Ibarra et al., 2012). DME targets gene-flanking small euchromatic transposons as well as many intergenic and heterochromatin sequences (Ibarra et al., 2012). How DME localizes to distinct targets with different chromatin states remains elusive, although the Facilitates Chromatin Transactions (FACT) histone chaperone is required for demethylation of heterochromatin and certain imprinted loci in the central cell (Ikeda et al., 2011;Frost et al., 2018). FACT complex is known to play a pivotal role in almost all chromatin-related processes, including transcription, replication, and DNA repair (Hondele and Ladurner, 2013). This is because nucleosomes are barriers to these processes that require access to the nucleosomal DNA, and the FACT complex is needed to facilitate destabilizing and disassembling nucleosomes. Interestingly, the FACT complex is only required for DME demethylation in central cell but not in vegetative nuclei (Frost et al., 2018), highlighting a difference in their chromatin conformation. There are three canonical histone H1 variants in Arabidopsis, the ubiquitous H1.1 and H1.2 variants and the stress-inducible H1.3 (Gantt and Lenvik, 1991;Ascenzi and Gantt, 1999;Wierzbicki and Jerzmanowski, 2005;Kotlinski et al., 2017). The vegetative cell has a decondensed nuclei and highly dispersed heterochromatins (Schoft et al., 2009); the two canonical linker histone H1.1 and H1.2 are specifically absent in the vegetative cell, which might contribute to the differential requirement of FACT for DME function (Hsieh et al., 2016;He et al., 2019).

H1 binds to the nucleosome core and the linker DNA between two adjacent nucleosomes and is required for higher-order chromatin organization such as heterochromatin compaction and chromocenter formation (Fyodorov et al., 2018). H1 loss causes dispersion of heterochromatin, nucleosome reorganization, and de-repression of H1-bound genes (Rutowicz et al., 2019;Choi et al., 2020). Since H1-bound nucleosomes are strong barriers to the DNA modifying enzymes such as DNA methyltransferases (Zemach et al., 2013;Lyons and Zilberman, 2017), loss of H1 facilitates DNA methyltransferases access to the decondensed heterochromatin and results in a gain in heterochromatic TEs methylation. However, euchromatic TEs exhibits a loss in DNA methylation in *h1* mutant, for reasons not fully understood (Zemach et al., 2013). Although earlier studies have shown that reducing the level of histone H1 affected the DNA methylation patterns of some imprinted genes in mammals and plants (Fan et al., 2005;Wierzbicki and Jerzmanowski, 2005;Rea et al., 2012), how H1 takes part in these processes has not been elucidated.

H1.2 interacts with DME in yeast two-hybrid *and in vitro* pull-down assays (Rea et al., 2012). In the endosperm derived from a cross between female *h1* triple mutant (*h1*.*1, h1*.*2-1, h1*.*3*) and wild-type pollen (referred to as *h1*-wt, or *h1* hybrid hereinafter), the expression of selected DME-regulated imprinted genes (*MEA, FIS2, FWA*) was reduced accompanied with an increase in the methylation of their maternal alleles, suggesting that H1 might play a role in mediating DME demethylation. However, loss of H1 differentially influences DNA methylation in heterochromatic and euchromatic TEs, raising the possibility that these effects might be epistatic to DME action. Here we showed that both *H1*.*1* and *H1*.*2* promoters are active in the central cell, with *H1*.*1* being more ubiquitously expressed in ovule while *H1*.*2* is specifically expressed in the central cell. Methylome analysis revealed that H1 loss decondensed the central cell nuclei and facilitated DME demethylation in heterochromatin, consistent with our observation that H1 is expressed in the central cell. Maternal H1 loss differentially affected methylation pattern of canonical DME target loci but did not substantially affect parent-of-origin specific expression of a list of reported imprinted genes. Our results revealed that loss of H1 indirectly affects DME-mediated DNA demethylation in the central cell and endosperm genome methylation but does not alter overall gene transcription or imprinting regulation in the endosperm.

## Materials and Methods

### Plant materials and growth conditions

*Arabidopsis Thaliana* plants were grown on soil with 3-day cold treatment before moving into growth chambers under 16 hour days/ 8 hour nights with temperatures of 23°C during the day and 22°C at night., Histone *h1* single T-DNA insertional lines were obtained from ABRC and used for generating *h1* double and triple homozygous mutants. The T-DNA line of *H1*.*1* is *h1*.*1-1* (SALK_128430C), which has a T-DNA insertion in the first exon, 133bp downstream of the start codon. The T-DNA line *h1*.*2-1* (Salk_002142) has the T-DNA insertion in the promoter region, 382bp upstream of the transcription start, and the T-DNA line *H1*.*3-1* (SALK_025209) has the T-DNA insertion also in the promoter region, 62bp upstream of the transcription start. We then searched for another *h1*.*2* T-DNA line to replace *h1*.*2-1* and found CS438975 (we named as *h1*.*2-2*), which has the T-DNA insertion in the 5’ UTR (untranslated region), 291bp downstream of the transcription start site. Histone *h1* double (*h1*.*1-1 h1*.*2-2* and *h1*.*2-2 h1*.*3-1*) and triple (*h1*.*1-1 h1*.*2-2 h1*.*3-1*) mutants were regenerated using *h1*.*2-2* T-DNA line. Expression of *H1*.*2* was almost undetectable in the *h1*.*2-2* mutant allele. All histone *h1*.*2*-containing mutants used in this research were from *h1*.*2-2*. For genetic crossing, *Arabidopsis* young flower buds at stage 13 were used as female plant for crossing. After the female plants were emasculated for 24 hours, pollen from several anthers were used to pollinate the same pistil to increase the chance of pollination. After pollination, the pistil was coved with a plastic bag to avoid contacting with other undesired pollen.

### GUS staining

The promoter region of histone *H1*.*1* was PCR amplified with primers H1.1promoter_HindIII_fwd and H1.1promoter_BamHI_rev. The promoter region of histone *H1*.*2* was PCR amplified with primers H1.2promoter_SalI_fwd2 and H1.2promoter_XbaI_rev2. The promoter region of histone *H1*.*3* was PCR amplified with primers H1.3promoter_HindIII_fwd and H1.3promoter_BamHI_rev. The amplified *H1* promoter regions were fused with reporter gene GUS (β-glucuronidase) in the binary vector pBI101, respectively. The constructs were confirmed by sequencing and transformed into *Arabidopsis* Col-0 by *Agrobacterium* infiltration. After screening for transgenic plants, stable transgenic T2 or later plants were used for GUS staining analysis. Tissue samples from young seedlings, different stages of floral buds and seeds were harvested and put immediately into the GUS staining buffer (10 mM sodium phosphate buffer, pH 7.2, 0.5% Triton X-100, and 1 mg/mL 5-bromo-4-chloro-3-indolyl-β-glucuronic acid) on ice. Samples in the staining buffer were vacuum infiltrated on ice for 30 minutes. Samples were incubated at 37°C with rotation for 18 hours or less. Tissues were destained using 30%, 50%, 70% ethanol in sequence for 30 minutes each time with rotation. Samples were imbibed in seed clearing solution or diH2O on slides for imaging.

### GFP expression

The *H1*.*1* promoter region was PCR amplified with primers H1.1promoter_HindIII_fwd and H1.1proSalIrev. The *H1*.*1* promoter region with gene body was PCR amplified with primers H1.1proSalIfwd and H1.1CDSBamHIrev. The *H1*.*2* promoter region was PCR amplified with primers his1.2 5’ SalI and his1.2 3’ XbalI. The *H1*.*3* promoter region was PCR amplified with primers H1.3promoter_HindIII_fwd and H1.3proSalIrev. The *H1*.*3* promoter region with gene body was PCR amplified with primers H1.3proSalIfwd and H1.3CDSBamHIrev. The amplified *H1* promoter regions or promoter regions with gene body were cloned into vector pBI-EGFP, respectively. The constructs were confirmed by sequencing and transformed into *Arabidopsis* Col-0 by *Agrobacterium* infiltration. After screening for transgenic plants, stable transgenic T2 or later plants were used for GFP expression analysis. Flowers before and after fertilization were harvested. Ovules at flower stage 12-13 and seeds were dissected out and imbibed in diH2O on slides for imaging using confocal microscopy. GFP was excited by the 488 nm laser line and was detected in a range between 505 nm to 535 nm.

The *pFWA*::GFP construct a gift from Dr. Tetsu Kinoshita (Yokohama City University) and was the same construct used in (Ikeda et al., 2011). The *MEDEA* promoter-GFP construct contains a 4.238 bp of MEA promoter (upstream of ATG), the HTB2-GFP (nuclear-localized GFP fused with the Arabidopsis histone H2B HTB2 (AT5G22880) protein), and a 2.1 kb *MEDEA* 3’- end sequence (downstream of the stop codon) concatenated using the Gibson assembly method. The backbone plasmid is a binary plasmid vector, pFGAMh (a hygromycin resistant version of pFGC5941) described before (Zhang et al., 2019).

### Purification of endosperm nuclei through fluorescence activated cell sorting (FASC)

Genetic crosses were made as described above. At 7 or 8 DAP (determined as the late heart or early heart stage embryo), embryo and endosperm were dissected out and collected under a dissecting microscope (Rea et al., 2012). For DNA extraction, collected embryo and endosperm were into 140ul nuclei extraction buffer with protease inhibitor as described (Zheng and Gehring, 2019); used pestle to grind material in the extraction buffer on ice. For each sample, grind about 2 min, then800ul nuclei staining buffer was added. Tubes containing nuclei staining buffer were wrapped withaluminum foil to avoid light. Filtered the nuclei in extraction buffer and staining buffer twice with CellTrics 30um filter on ice. Then kept filtered nuclei on ice. The nuclei were sorted at the Flow Cytometry Research Core Facility Doisy Research Center (St. Louis University) with a BD FACSAriaI IIu equipped with a 407nm violet laser using the following parameters: Nozzle size: 100 microns; Sheath pressure: 30 psi; Droplet frequency: 28,100 drops/sec.; Precision mode: Purity. Nuclei were gated based on signals from the DAPI channel.

For RNA extraction, embryos and endosperms were put into 1.5ml empty Eppendorf tubes floated on liquid nitrogen. After collection, embryos and endosperms are kept in -80C. Nuclei extraction buffer and nuclei staining buffer are added later before cell sorting. After sorting, Purified embryo and endosperm were collected with 1.5ml Eppendorf tube with 50 μl TRIzol™ Reagent (Invitrogen), Zymo RNA Lysis buffer or other RNA lysis buffer which were ready for RNA extraction.

### Whole-genome bisulfite sequencing and DNA methylome analysis

Crosses were performed between *h1* triple mutant *h1*.*1-1 h1*.*2-2 h1*.*3-1* (female plant) and wild type L*er* (male plant). Embryos and endosperms were collected 8 days after crossing and purified through FACS. Genomic DNA was isolated from FACS-purified nuclei using CTAB method as described (Ibarra et al., 2012). Approximately 2-5 ng of purified genomic DNA was spiked with 1% (w/w) of unmethylated cl857 *Sam7* Lambda DNA (Promega, Madison, WI) and sheared to about 400bp using Covaris M220 (Covaris Inc., Woburn, Massachusetts) under the following settings: target BP, 350; peak incident power, 75 W; duty factor, 10%; cycles per burst, 200; treatment time, 60 second; sample volume 50μl. The sheared DNA was cleaned up and recovered by 1.2x AMPure XP beads followed one round of sodium bisulfite conversion using the EZ DNA Methylation-Lightning Kit (Zymo Research Corporation, Irvine, CA) as outlined in the manufacturer’s instruction with 80 min of conversion time. Bisulfite sequencing libraries were constructed using the ACCEL-NGS Methyl-Seq DNA library kit (Product Code 30024, Swift Biosciences, Ann Arbor, MI) per the manufacturer’s instruction. The PCR enriched libraries were purified twice with 0.8x (v/v) AMPure XP beads to remove adaptor dimers. High throughput sequencing was performed by Novogene Corporation (Davis, CA.). Sequencing reads from three individual transgenic lines were used in the analysis (Table S9). Sequenced reads were mapped to the TAIR10 (whole genome) and TAIR8 (allele-specific) reference genomes and DNA methylation analyses were performed as previously described (Ibarra et al., 2012).

Fractional CG methylation in 50-bp windows across the genome was compared between embryo and endosperm from *h2ax* seeds as well as wild-type embryo, endosperm, and *dme-2* endosperm (GSE38935). Windows with a fractional CG methylation difference of at least 0.3 in the endosperm comparison (Fisher’s exact test p-value < 0.001) were merged to generate larger differentially methylated regions (DMRs) if they occurred within 300 bp. DMRs were retained for further analysis if the fractional CG methylation difference across the merged DMR > 0.3 (Fisher’s exact test *p-*value < 10-6), and if the DMR is at least 100-bp long. The merged DMR list is in the supplemental Dataset S1. Distribution of DMRs along genes and whole genome methylation average metaplots of Genes and TEs were plotted as described previously (Ibarra et al., 2012;Zhang et al., 2019).

### RNA sequencing

Crosses were performed between *h1* triple mutant *h1*.*1-1 h1*.*2-2 h1*.*3-1* and wild type L*er*. Crosses between Col-0 and L*er* were used as the control. Embryos and endosperms were collected 8 days after cross and total RNA was extracted using RNAeasy kit (Qiagen) plus in-column DNase digestion. Three independent sets of total RNAs from embryo and endosperm of *h1* Col x L*er* and wild-type Col x L*er* were isolated. Illumina cDNA libraries were constructed with the Ovation RNA-seq System V2 (NuGen Technologies) per manufacturer’s instruction and as described (Hsieh et al., 2011). Adapter trimming was performed using the fastp program (Chen et al., 2018). For allele-specific transcriptome analysis, adapter-trimmed sequencing reads were mapped to the TAIR8 Col-0 and L*er* cDNA scaffold using custom scripts as before (Hsieh et al., 2011). Individual reads were assigned to Col-0 or L*er* ecotype based on the SNPs detected between the two ecotypes. Each gene received Col-0 and L*er* scores, which represented the number of reads aligned to the corresponding ecotype. These numbers were used to calculate the maternal transcript proportion for each gene. For endosperm and embryo transcriptomes, adapter-trimmed sequencing reads were aligned to TAIR10 for transcript assignation and quantification with Salmon (v.1.9.0) (Patro et al., 2017). DESeq2 was used to identify differentially expressed genes (Love et al., 2014).

## Results

### Histone *H1*.*2* is expressed in the central cell of female gametophyte and developing endosperm

Although DME is preferentially active in the central cell, it is also expressed in the pollen vegetative cell (VC) and demethylates the VC genome during *Arabidopsis* reproduction (Schoft et al., 2011;Ibarra et al., 2012). However, H1.1 and H1.2 are present in sperm but undetectable in vegetative nuclei and H1.3 expression is not detected in pollen (Hsieh et al., 2016;He et al., 2019). This indicates that H1 is not required for DME demethylation in VCs. To examine whether histone *H1* is expressed in central cell, we fused three *H1* promoter regions with the *GFP* reporter gene respectively and transferred these GFP fusion constructs into *Arabidopsis* Col-0. Although both *H1*.*1* and *H1*.*2* were reported to be widely expressed (Rutowicz et al., 2015), we were surprised to see that *H1*.*2* was only expressed in the central cell of stage 12 flowers but undetectable during earlier gametogenesis (**Figure 1**). After fertilization, *H1*.*2* expression is observed in the nuclear cytoplasmic domain (NCD) of early endosperm, but the expression level decreases as endosperm develops (**Figure 1**). *H1*.*2* expression was not observed in embryo or any part of the female gametophyte other than the central cell whereas the *H1*.*1* promoter-GFP construct was more ubiquitously expressed in ovules and seeds including the integument, central cell, embryo, and endosperm (**Figure S1**). Using the same set of *H1* promoters driving the GUS reporter gene confirmed what was reported that both *H1*.*1* and *H1*.*2* are widely expressed, with *H1*.*1* being more abundantly expressed than *H1*.*2* whereas *H1*.*3* is very lowly expressed in the tissues assessed (**Figure S2**). Supporting our observation of central cell-specific *H1*.*2* expression, Song et al reported that *H1*.*2* had an average expression of 135.86 (TPM normalized) in the central cell which is about 15-fold higher than the average expression of 9.26 in the egg cell (Song et al., 2020). The Evorepro database (https://evorepro.sbs.ntu.edu.sg/) also revealed that *H1*.*2* expression in ovules is more than 4-fold that in the egg cell (**Figure S3**), which is consistent with our result.

**FIGURE 1.**
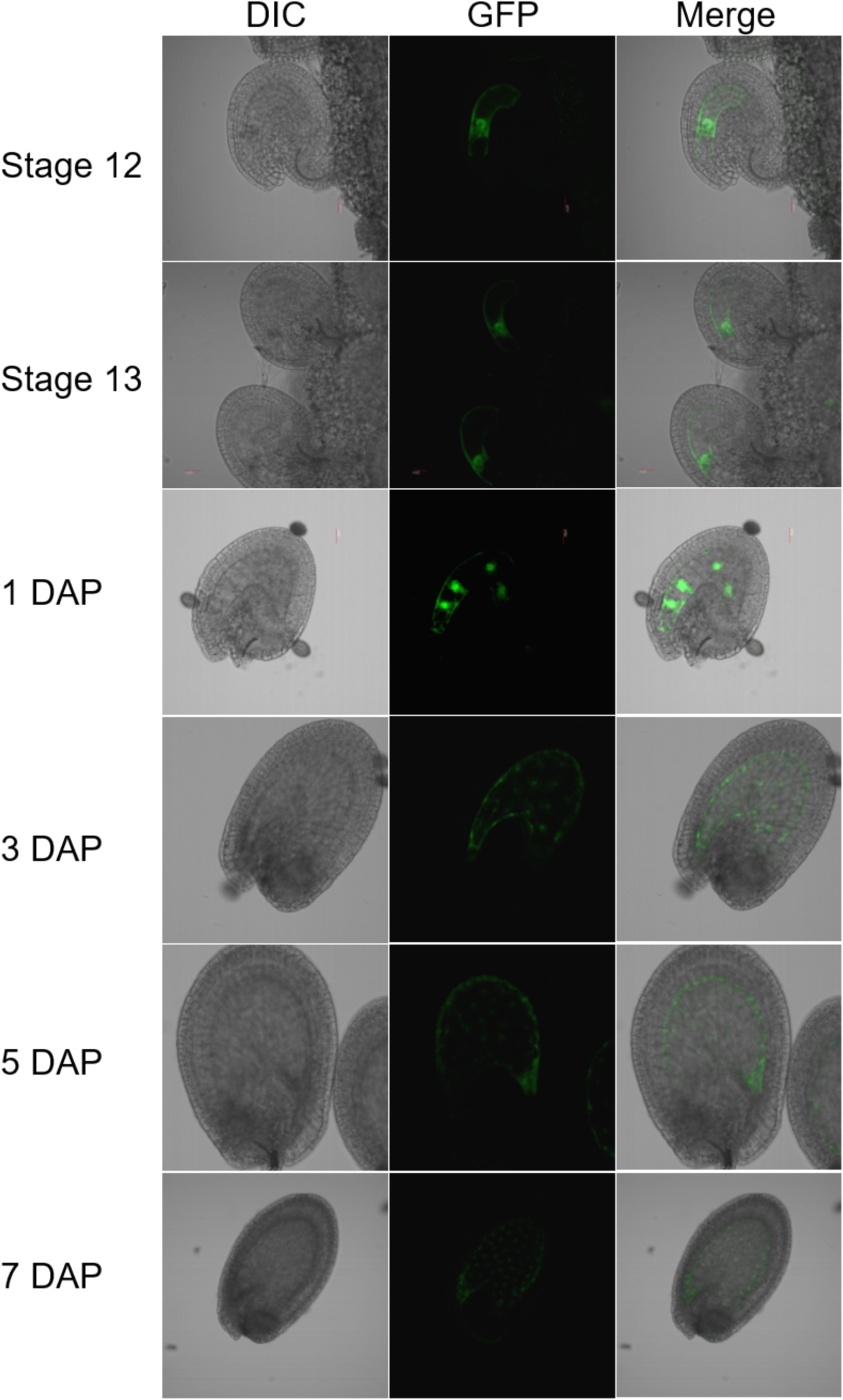
Expression of the *H1*.*2 promoter::GFP* transgene in *Arabidopsis* ovules and young developing seeds. Ovules and seeds with expression of *H1*.*2 promoter*:*GFP* transgene were photographed using confocal fluorescence microscope. Ovules at flower stage 12 and stage 13 and seeds at 1, 3, 5, and 7 DAP were hand-dissected for imaging. The GFP signal is shown in green. DAP, Days After Pollination; DIC, Differential Interference Contrast.

Central cell-specific expression of *H1*.*2* was unexpected. We wondered whether *H1*.*2* expression in the central cell is regulated by *DME*. To test this, we crossed the homozygous *pH1*.*2:GFP* transgenic plants with *DME/dme-2* heterozygotes and obtained the F1 plants that were heterozygous for *DME/dme-2* and hemizygous for the *pH1*.*2:GFP* transgene. In the F2 generation, we examined the *pH1*.*2:GFP* expression in the *DME/dme-2* mutant and their segregating wild-type siblings. We observed that 48% of the female gametophytes with ovules showed strong GFP signal in the *DME/DME pH1*.*2:GFP*/-(87 ovules expressed GFP out of 183 total ovules, 87:96, 1:1, χ2 = 0.44, P > 0.50). and 47% of expression in the *DME/dme*-2 *pH1*.*2:GFP*/-plants (83 ovules expressed GFP out of 176 total ovules, 83:93, 1:1, χ2 = 0.56, P > 0.46). This result indicates that *H1*.*2* expression in the central cell is not regulated by *DME*. This is consistent with *H1*.*2* not being an imprinted gene as we reported earlier (Rea et al., 2012).

### Loss of histone H1 affects the expression of *MEA* and *FWA* before and after fertilization

We previously showed that *h1*-wt hybrid endosperm (*h1* x L*er*) reduced the expression of *MEA* and *FWA* (Rea et al., 2012). To gain a better understanding on how loss of H1 affects *MEA* and *FWA* expression, we used *MEA* and *FWA* promoter::*GFP* constructs and transferred them into *Arabidopsis* wild type Col-0 and the *h1* triple mutant. Transgenic plants homozygous for *pMEA:GFP* and *pFWA:GFP* were used to examine *MEA* and *FWA* gene expression. In wild-type plants, *MEA* is specifically expressed in the central cell nucleus of stage 12 flower and in the endosperm after fertilization (**Figure 2**). In *h1* mutant, fluorescent intensity of the *pMEA:GFP* transgene was much lower than that in the wild type and the number of GFP positive ovules and seeds was significantly reduced. We detected 67.1% of *h1* ovules (94 out of 140) expressing the *pMEA:GFP* transgene compared with 85.3% (262 out of 307) in wild-type at stage 12 flowers (**Table 1**). As the seeds develop, the number of *h1* mutant seeds expressing the *pMEA:GFP* transgene increased to 79.1% compared to 91.5% in WT at 1 day after pollination (DAP) and reached 94.4% at 4 DAP which was comparable with what was seen in wild-type (95.2%). However, the GFP fluorescent intensity was noticeably lower in the *h1* mutant than in wild type using the same parameter settings under the fluorescence microscope. The *pFWA:GFP* transgene also exhibited a similar expression pattern as the *pMEA:GFP* transgene. In the *h1* triple mutant seeds at 1 DAP, the GFP fluorescent intensity was much lower than that in wild type and the percentage of *pFWA:GFP* expression seeds (84.5%) was less than wild type (94.6%) (**Figure S4, Table 1**). As the *h1* triple mutant seeds developed, nearly all seeds expressed the *pFWA:GFP* transgene, but the GFP intensity remains lower than in wild type. In summary, *h1* mutations reduce *MEA* and *FWA* gene expression in the central cell and endosperm.

**FIGURE 2.**
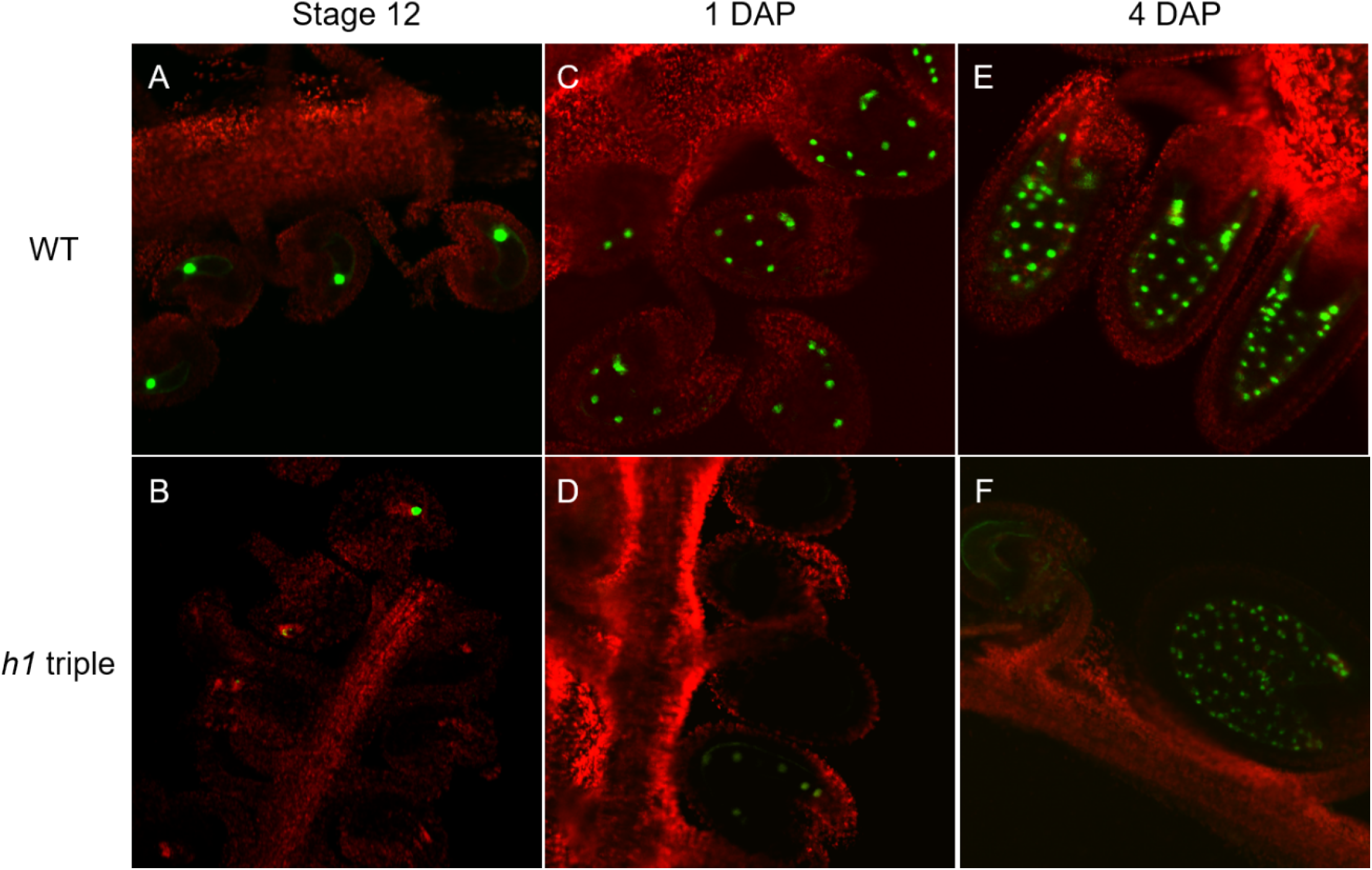
Expression of the *H1*.*2 promoter::GFP* transgene is reduced in the central cell of *h1* hybrid endosperm. Fluorescence images of pMEA::GFP expression signals in wild type (A, C, E) and in h1 triple mutant endosperm (B, D, F). The GFP and chlorophyll fluorescence signals were pseudo-colored as green and red, respectively. Fluorescence micrographs of ovules at flower stage 12 (A, B), 24-hr after pollination (C, D), and 96-hr after pollination (E, F).

**Table 1.**
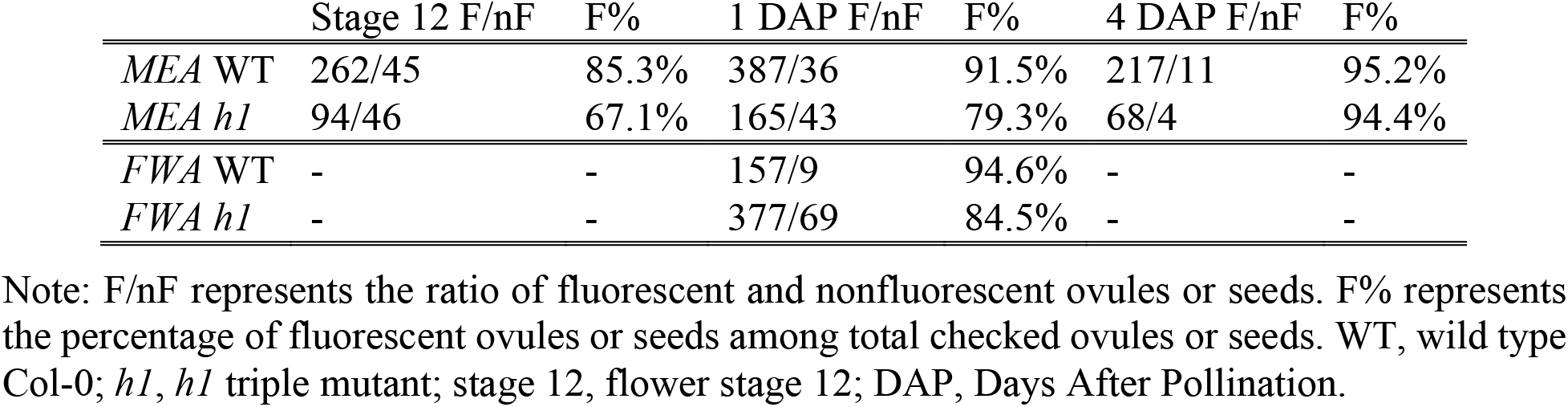
The effect of the *h1* mutation on the expression of *MEA* and *FWA* in the central cell and endosperm.

### Maternal H1 loss influences DME-mediated DNA demethylation in the central cell and endosperm genome methylation

The reduction in *MEA* and *FWA* expression and the increase in DNA methylation in their maternal alleles in *h1*-wt hybrid endosperm (Rea et al., 2012) suggested that H1 might play a role in assisting DME demethylation in the central cell. If H1 is necessary for DME demethylation, the majority of canonical DME targets would exhibit a loss of DNA demethylation (or hypermethylation) in the hybrid endosperm. Alternatively, the reduced *MEA* and *FWA* expression seen in the hybrid endosperm could be due to a locus-specific effect caused by maternal *h1* mutations that makes it less favorable for DME demethylation. In this scenario, we would expect to see a majority of DME target sites are properly demethylated accompanied by specific loci exhibiting an increased or reduced level of DNA methylation. Since loss of DME results in a striking seed abortion phenotype (Choi et al., 2002), the largely normal *h1* mutant seeds (Rea et al., 2012) suggested that DME demethylation should be largely intact in these seeds, at least in the DME-regulated Polycomb Repressive Complex 2 (PRC2) genes (*i*.*e*., *MEA* and *FIS2*) critical for seed viability (Choi et al., 2002;Jullien et al., 2006). To investigate the effect of *h1* mutations on DME demethylation, we carried out genome-wide bisulfite sequencing of the F1 hybrid endosperm derived from crosses between *h1* triple mutant or wild-type female plants (Col-0) and wild-type L*er* as male parents. To avoid contamination of seed coat tissues, we prepared crude nuclei from the hand dissected embryo and endosperm tissues from the hybrid seeds and used fluorescence activated cell sorting (FACS) to isolate pure embryo (2C and 4C) and endosperm (3C and 6C) nuclei for BS-seq library construction and sequencing (**Figure S5**). We used the Swift Bioscience’s Adapase™ technique (see Materials and Methods) that was successfully used for methylation profiling from ultra-low amounts of input genomic DNA (Luo et al., 2017;Luo et al., 2018). Methylomes from three biological replicates of *h1* x L*er* embryo and endosperm were generated. We used the Col-L*er* SNPs to sort and assign uniquely mapped reads to their respective parents of origin. The maternal:paternal read ratios for endosperm and embryo libraries tightly followed the expected ratios (2:1 for endosperm and 1:1 for embryo), indicating sample purity of FAC sorted nuclei with little maternal seedcoat contamination (**Table S1**). Pearson correlation coefficients were highly concordant between bio-reps (**Table S2**) and reads from the same tissue/genotype were pooled for subsequent analysis. We first looked at the methylation profiles of selected DME target genes in genome browser. Their methylation profiles are clearly lower in the histone *h1* hybrid endosperm than that in the histone *h1* hybrid embryo. However, the methylation levels of *FWA* and *MEA* were higher in *h1* hybrid endosperm compared with wild type (**Figure S6**, top panel), which is consistent what we reported before that demethylation of the maternal alleles in these loci were less effective in the *h1* endosperm (Rea et al., 2012). By contrast, *YUC10, SDC*, and *DRB2* exhibited a lower DNA methylation profiles in the *h1* hybrid endosperm (**Figure S6**, lower panel), suggesting that maternal *h1* mutations variably influenced the methylation of certain imprinted genes.

The two polymorphic parental strains (Col x L*er*) allowed assessment of methylation difference between the two parental genomes. In wild-type endosperm, kernel density plot of fractional DNA methylation difference between the paternal L*er* and the maternal Col genome exhibited a center peak around zero and a small population of 50-bp windows toward the positive tail-end, representing localized demethylated regions hypomethylated on the maternal genome by DME (Ibarra et al., 2012) (**Figure S7A**, blue trace). The fractional methylation difference between paternal (wild-type L*er*) and maternal (*h1* Col) genomes also displayed a zero-centered peak and a more profound positive end bump, suggesting a more extended demethylation on the maternal *h1* genome (**Figure S7A**, red trace). However, the availability of SNPs between the Col-0 and L*er* genomes (∼400,000 SNPs) (Hsieh et al., 2011;Ibarra et al., 2012) limits assessment of only a small fraction of the genome, we instead focused subsequent analysis at the genome-wide scale.

Wild-type endosperm and embryo methylome comparison is one accepted proxy for the comparison of DME actions in the central cell versus the egg cell because DME is only active in the central cell but not in the egg cell of mature female gametophyte (Hsieh et al., 2009). We reasoned by comparing the differentially methylated regions between wild-type embryo versus endosperm (referred to as wt-DMRs, as the proxy for DME activity in wild-type central cell) and the DMRs between of the histone *h1* mutant embryo versus endosperm (*i*.*e*., *h1*-DMRs, as the proxy for DME activity in the *h1* mutant central cell) we can gain valuable insights into how loss of H1 affects DME demethylation. Since the canonical DME target sites are well-defined (Ibarra et al., 2012), we separated the genome into DME and non-DME target sites and used kernel density estimate to compare the embryo-endosperm difference in wild type versus and in *h1* hybrid seeds. For the DME sites, embryo and endosperm CG methylation difference in wild-type (**Figure 3A**, blue trace) and *h1* hybrid seeds (green trace) nearly overlap, indicating that the DME action is largely intact without H1. For regions outside canonical DME sites, the embryo and endosperm difference are reduced (**Figure 3A**, orange trace, peak shifted toward zero) in *h1* hybrid seeds compared with wild type (**Figure 3A**, red trace, peak is slightly toward the positive side). Since CG methylation in embryo is higher than in endosperm (Hsieh et al., 2009;Ibarra et al., 2012), this reduction indicates a slight overall CG hypermethylation of non-DME target sites in the *h1*-wt hybrid endosperm. These features suggest that there is no large-scale change in DNA methylation profile in *h1*-wt hybrid seeds.

**FIGURE 3.**
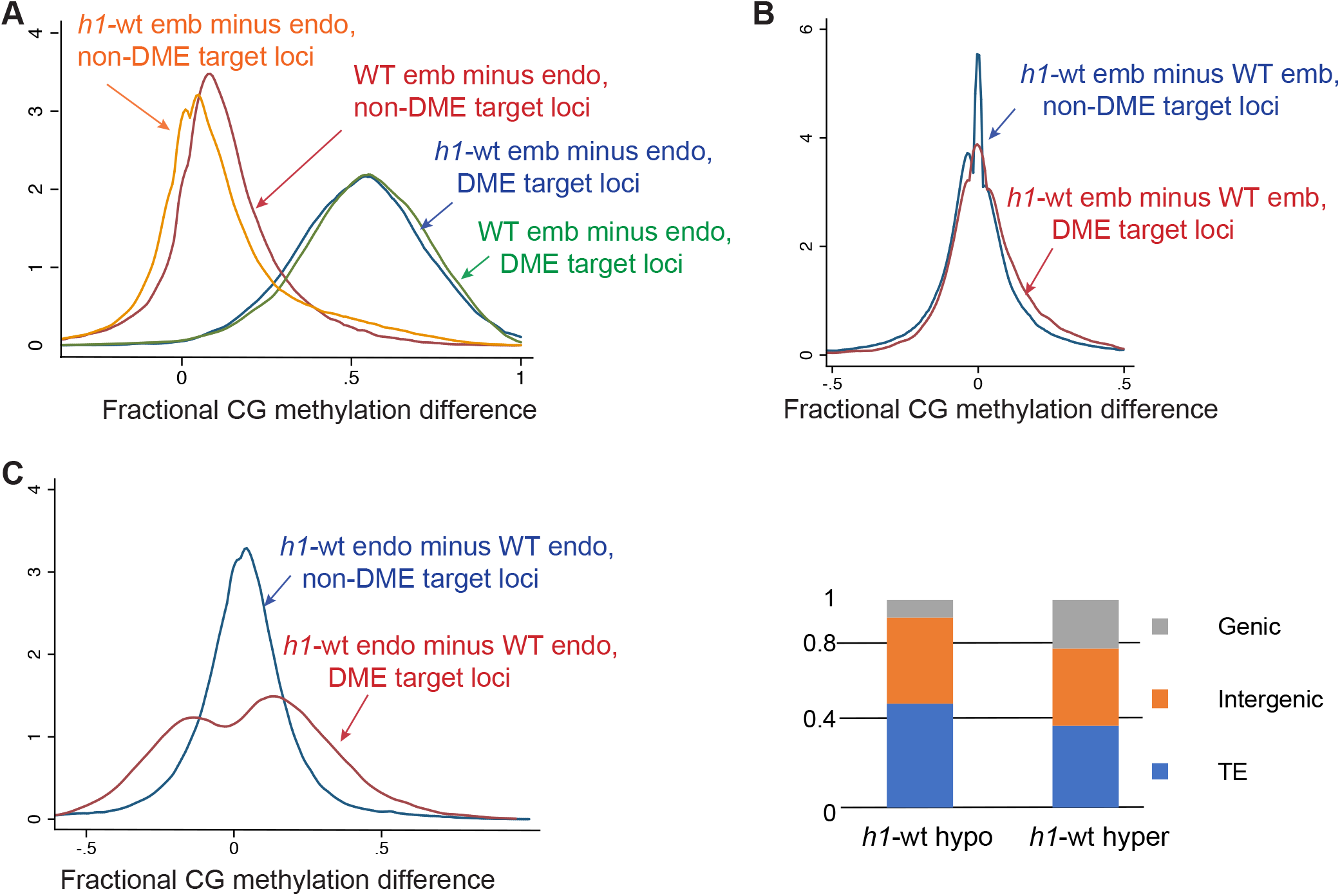
Methylome analysis of embryo and endosperm derived from the maternal wild type or *h1* mutant crossed with L*er* pollen. (A) Kernel density plots of CG methylation difference between embryo and endosperm (embryo minus endosperm) in wild type Col x L*er* (blue and magenta traces) or in hybrid *h1* x L*er* endosperm (green and orange traces). Methylated loci were grouped into canonical DME targets (blue and green traces) and non-target loci (magenta and orange traces). (B) Kernel density plots of CG methylation difference between wild-type and hybrid *h1* embryo in DME targets (magenta) and non-targets (blue). (C) Kernel density plots of CG methylation difference between wild-type and hybrid *h1* endosperm in DME targets (magenta) and non-targets (blue). (D) Percent distribution of the three different genomic features in canonical DME targets that loose (*h1*-hypo) or gain (*h1*-hyper) in hybrid *h1* endosperm compared with wild type.

We next compared the difference between *h1*-wt hybrid and wild-type methylomes. DNA methylation was largely unchanged between in the *h1*-wt and wild-type embryos with the peak of fractional methylation difference sharply centered at zero (**Figure S7B**, blue trace) franked by two minor shoulder peaks, indicating that localized minor hyper and hypo methylations exist in the mutant embryo. This is also evident in whole genome methylation metaplots showing near identical CG methylation profiles between wild type and *h1* hybrid embryos (**Figure S8**). By contrast, CG DNA methylation of the *h1*-wt endosperm is higher compared to wild-type endosperm, with the fractional methylation difference peak shifted toward the right side and flanked by broader shoulders (**Figure S7B**, red trace). This suggests that a slight CG hypermethylation occurs in the *h1*-wt endosperm. Whole genome CG metaplots revealed this hypermethylation occurs in coding sequences and to a higher degree in longer TEs (**Figure S8**). This likely reflects earlier reports that heterochromatin TEs gain DNA methylation in *h1* mutant due to easier access for DNA methyltransferase to the DNA templates (Lyons and Zilberman, 2017) (Zemach et al., 2013). Interestingly, this heterochromatin TE hypermethylation is not observed in the *h1* embryo as little change in DNA methylation is observed between *h1*-wt and wild-type embryos (**Figures 3B and S8**).

To gain a better insight into how H1 loss affects DME demethylation, we plotted the fractional methylation difference between *h1*-wt and wild type within and outside of the canonical DME target sites. Whereas very little difference between these two groups of loci was observed in embryo (**Figure 3B**), a more widespread difference can be seen within the canonical DME target sites in the endosperm (**Figure 3C**, red trace), indicating that the loss of H1 differentially affected demethylation at the canonical DME target loci. DME targets more demethylated in *h1*-wt endosperm (**Figure 3C**, *h1*-hypo loci, fractional methylation difference <0, left peak of red trace) are enriched for heterochromatin (50.2% versus 39.5%) but depleted for genic sequence (8.2% versus 23.6%) (**Figure 3D**) relative to DME targets hypermethylated in the *h1*-wt endosperm (**Figure 3C**, *h1*-hyper loci, difference >0, right peak of red trace). This suggest that without H1, certain heterochromatin loci are more demethylated, which is consistent with the model that H1-containing nucleosomes restrict accessibility of chromatin modifying enzymes (Zemach et al., 2013;Lyons and Zilberman, 2017). Why some DME loci (**Figure 3C**, *h1*-hyper sites) are less demethylated in the *h1*-wt endosperm is unknown, but they are enriched for chromatin states 3 and 7, hallmark features for intragenic sequences (**Figure S9**) (Sequeira-Mendes et al., 2014). For loci outside of canonical DME targets, there is also a slight difference between *h1* and wild-type endosperm (**Figure 3C**, blue trace with peak shifted toward positive side and a broader shoulder compared with embryo plot). Thus, the *h1* mutant specifically affects the methylation pattern of hybrid endosperm but not embryo, implicating the presence of an altered DME demethylation activity in the *h1* mutant central cell.

We next compared the differentially methylated regions (DMRs) between wild-type embryo versus endosperm (DME targets in the presence of H1 or WT-DMRs, n=8207), and the DMRs between *h1*-wt embryo versus endosperm (DME targets in the absence of H1 or *h1*-DMRs, n=11552) (Materials and Methods) using previously established criteria (supplementary dataset 1, DMR lists) (Ibarra et al., 2012). The *h1*-DMRs cover over 6 million bases, more than twice longer than the WT-DMRs (**Figure 4A**). About 56% of the *h1*-DMRs overlap with WT-DMRs (**Figure 4B**). On average, the *h1*-DMRs are significantly longer in size (**Figure 4C**, t=29.1221, *p*=0, Welch’s t-test). These observations suggest that without H1, DME demethylation is less constrained. Natural depletion of H1 in the Arabidopsis vegetative cell (VC) is associated with heterochromatin de-condensation and ectopic H1 expression in VC impedes DME from accessing heterochromatic transposons (He et al., 2019). Consistent with this model, the *h1*-unique DMRs are highly enriched for the heterochromatin states (State 8 & 9, **Figure 4D**) (Sequeira-Mendes et al., 2014). This is also supported by the DMR distribution plot across the genome showing *h1*-DMRs are more abundant in the pericentromeric and genic regions (**Figure 4E, 4F**).

**FIGURE 4.**
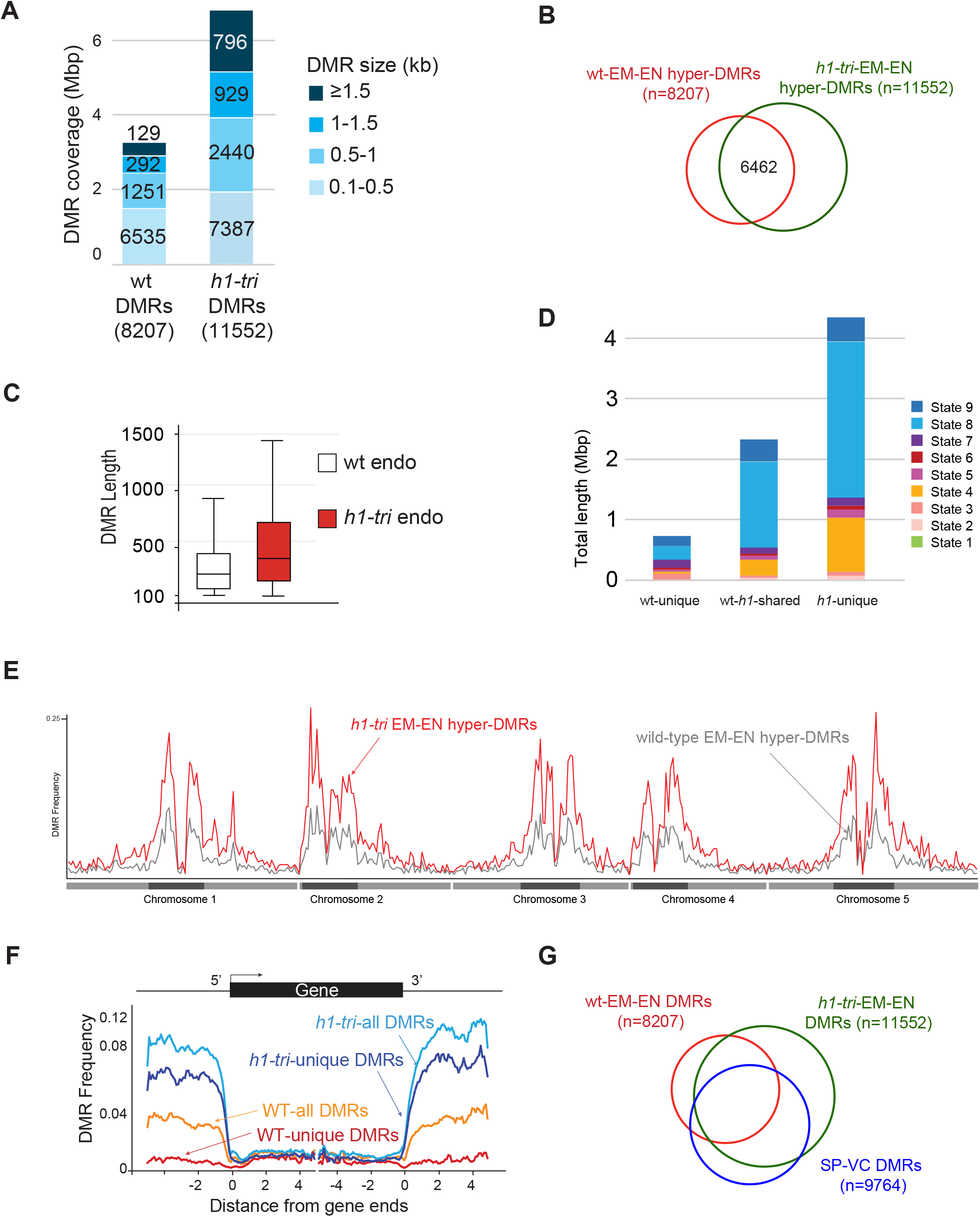
Analysis of the hyper DMRs of embryo vs endosperm derived from maternal wild type or *h1* mutant crossed with L*er* pollen. (A) Wild-type and *h1* DMRs grouped by size, with the cumulative total length they cover shown. (B) Venn diagram depicting overlaps between Wild-type and *h1* DMRs. (C) Boxplot showing the length distribution of Wild-type and *h1* DMRs. (D) Chromatin state distribution, and the total length they covered, within wt-unique, wt-*h1* shared, and *h1*-unique DMRs. States 1 to 7 correspond to euchromatin, and states 8 and 9 correspond to AT- and GC-rich heterochromatin, respectively. (E) Distribution frequency of DMRs along the 5 chromosomes. Dark blocks represent centromere and peri-centromeric regions of each chromosome. (F) Distribution frequency of DMRs with respect to coding genes. Genes were aligned at the 5’- or the 3’-end, and the proportion of genes with DMRs in each 100-bp interval is plotted. DMR distribution is shown with respect to all wt-DMRs (orange trace), wt-unique DMRs (red trace), all *h1*-DMRs (light blue trace), and *h1*-unique DMRs (dark blue trace). (G) Venn diagram showing overlaps between Wild type, *h1*, and sperm vs vegetative cell DMRs.

DME is also active in the vegetative cell where it demethylates a slightly larger number of loci (referred to as SP-VC DMRs, n=9764) than in the central cell (canonical DMRs, *dme* mutant versus wild-type endosperm, n=8672) (Ibarra et al., 2012). About 56% of the DME targets in the female gametophyte (canonical DME DMRs) and 50% of WT-DMRs (EM vs EN) overlap with the SP-VC DMRs (Ibarra et al., 2012). One factor attributed to this moderate overlap between the male and female gametophytes is the H1-depleted, heterochromatin-decondensed vegetative nuclei that is thought to facilitate DME access to the heterochromatin targets (He et al., 2019). Notably, greater than 80% (and up to 86%) of known DME DMRs (i.e., the SP-VC DMRs, canonical DME DMRs, and WT-DMRs) overlap with the *h1*-DMRs (**Figure 4G, Table S3**). These observations strongly support the model that lack of H1 facilitates DME access to the heterochromatin regions, which was the primary attribute for the increased number of *h1*-DMRs.

### Maternal *h1* mutations do not alter gene transcription and imprinting regulation in Arabidopsis endosperm

To assess how loss of H1 influences endosperm gene transcription and imprinting regulation, we manually dissected and collected endosperm from 7-DAP hybrid F1 seeds derived from Col-0 x L*er* and *h1* (Col-0) x L*er* crosses. Since endosperm RNA-seq using manually dissected materials is prone to seed coat contamination (Schon and Nodine, 2017), we first tried to isolate nuclei RNAs from FACS-purified 3C and 6C endosperm nuclei. However, due to the low input amount of dissected endosperm and instability of nascent transcripts in the nuclei, we were unable to purify any detectable amounts of nuclei RNAs in our experimental setting using multiple RNA-isolation methods. We therefore proceeded with regular RNA-seq analysis using manually dissected endosperm (Hsieh et al., 2011). We used the tissue enrichment test tool (Schon and Nodine, 2017) to evaluate the extend of seedcoat contamination in our endosperm and embryo RNA-seq datasets and found that indeed a modest degree of seedcoat enrichment can be detected, with a general seed coat (GSC) enrichment score of 5 to 10 among endosperm samples and 1.8 to 3 among embryo datasets (**Figure S10**), which is consistent with, but slightly lower than, all the manually dissected endosperm datasets assessed by an earlier study (Schon and Nodine, 2017). Pearson correlation coefficients between biological replicates showed that they were highly concordant (**Table S4**). Principal component analysis showed that samples were primarily separated by tissue type, with the PC1 capturing 82% of variation while genotype difference does not differentiate tissue samples (**Figure S11**). This is also reflected in the low numbers of up- and down-regulated DEGs between *h1* and wild-type endosperm and embryo (fold change >2, adjusted *p* value < 0.05, 31 up and 20 down in endosperm, 5 up and 13 down in embryo, **supplemental dataset 3**). No GO term enrichment was detected, likely due to the small numbers of DEGs identified. RNA-seq data showed that the expression of *FIS2* and *FWA* were reduced in the *h1*-wt endosperm (with a log2 fold change of -0.41 for *FIS2* and -1.23 for *FWA*), which is consistent with what we had observed before (Rea et al., 2012). The expression of *MEA* varied between replicates but overall unchanged between wild-type and *h1* endosperm (log2 fold change 0.2). All three genes were expressed at low levels. Since histone *H1* genes are not imprinted in the endosperm (Rea et al., 2012), loss of H1 function in the gamete and central cell is likely compensated by the functional paternal copies upon fertilization. By contrast, a complete loss of H1 in the homozygous *h1* triple mutant caused a significant mis-regulation of > 900 genes in leaves and seedlings (Choi et al., 2020) and 701 gene in Arabidopsis seedlings (Rutowicz et al., 2019). Maternal depletion of H1 also did not cause TE mis-regulation (3 up- and 2 down-regulated in endosperm, 2 up- and 4 down-regulated in embryo), in contrast to the 18 TEs significantly dysregulated in *h1* plants (Choi et al., 2020).

We used the Col-L*er* SNPs to distinguish and sum the number of reads derived from their respective parents as we did before (Hsieh et al., 2011). To minimize mis-interpretation of possible seedcoat contamination that could skew the maternal ratio of expressed genes, we focused our analysis only on a published list of high confident imprinted gens (148 MEGs and 81 PEGs) (Schon and Nodine, 2017) and assessed any difference between wild-type and *h1*-wt endosperm data that were generated by the same procedure. Among them, 80 MEGs and 42 PEGs have sufficient read numbers for proper comparison among our datasets. Overall, that maternal ratios of MEGs and PEGs are similar between wild-type and *h1* endosperm, with the MEGs showing a small increase in the maternal ratio (student’s t-test, P < 0.01) but not the PEGs (P > 0.05) (**Figure 5A**). Scatter plot of maternal transcript proportions among analyzed imprinted genes showed that they are highly correlated between wild-type and *h1*-wt endosperm (Pearson correlation r = 0.91, n=122, **Figure 5B**) indicating that loss of maternal H1 activity does not affect overall gene imprinting regulation. Genome-wide the correlation of maternal transcript proportion between wild-type and *h1* endosperm is also high (r = 0.83, n=9054, **Figure S12**), consistent with the low DEGs detected between them. In summary, our results show that maternal genome *h1* mutations have a very limited effect on endosperm gene transcription or imprinted regulation.

**FIGURE 5.**
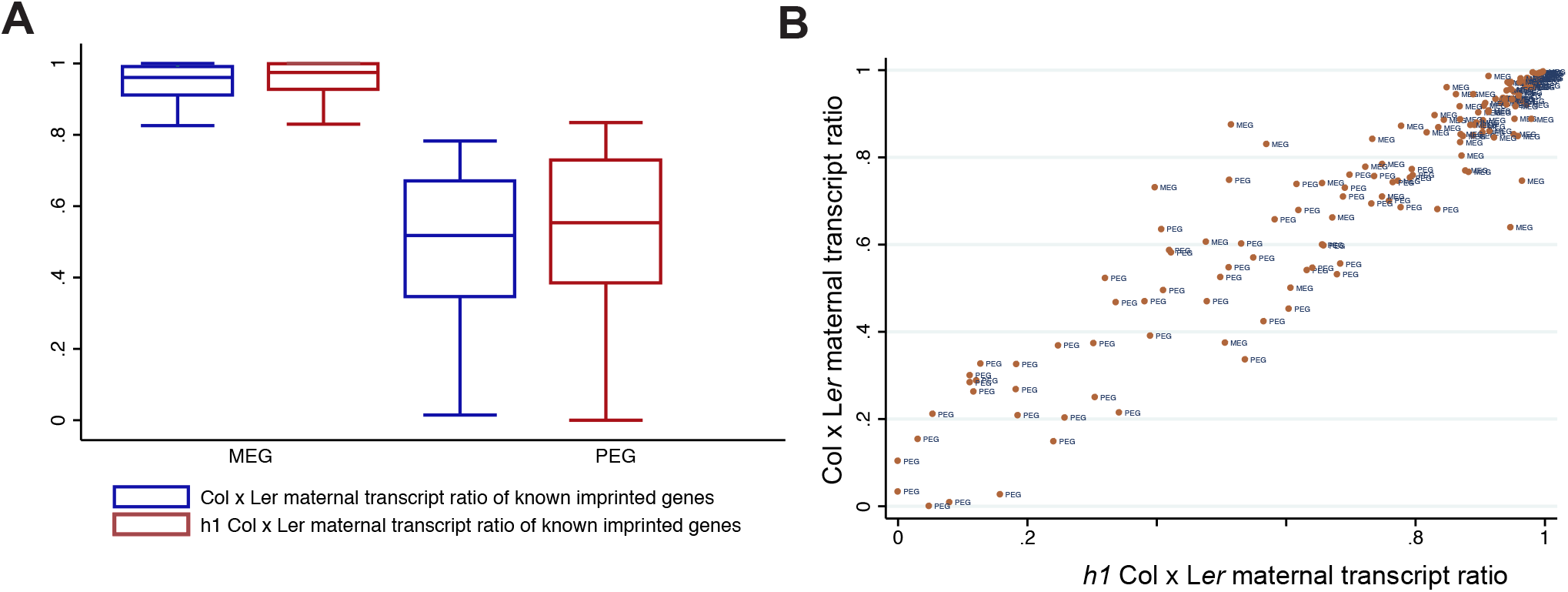
Known imprinted genes and their status of imprinted expression in wild type and *h1*- wt hybrid endosperm. (A) Boxplot showing the distribution of maternal transcript proportions of selected MEGs and PEGs in wild type and h1-wt hybrid endosperm. (B) Scatterplot showing the correlation of maternal transcript proportion of each imprinted gene between wild type and *h1*-wt hybrid endosperm.

## DISCUSSION

DME, a multi-domain novel glycosylase that initiates active DNA demethylation process by removing the methylated cytosine bases in the central cell, regulates gene imprinting and is essential for seed development in *Arabidopsis* (Choi et al., 2002) (Ibarra et al., 2012) (Gehring et al., 2006). In pollen, DME also demethylates the VC genome at thousands of loci to reinforce gamete TE silencing and to ensure robust pollen germination in certain accessions (Schoft et al., 2011;Ibarra et al., 2012). DME preferentially targets small and genic-flanking transposons whose demethylation influences the transcription of nearby coding genes, it also demethylates many intergenic and heterochromatin targets. How DME is specifically recruited to its target loci remains elusive, due to its ephemeral expression in the gamete companion cells not accessible by common experimental methods. Using a yeast two-hybrid screen, Rea et al. identified linker histone H1.2 as a candidate DME-interacting protein and demonstrated that they did physically interact in an *in vitro* pull-down assay (Rea et al., 2012). Furthermore, in the female *h1* triple mutant (*h1*.*1, h1*.*2-1, h1*.*3*) x wild-type L*er* endosperm, the expression of selected DME-regulated imprinted genes (*MEA, FIS2, FWA*) were reduced and the methylation of their maternal alleles were increased, suggesting that DME demethylation might be impaired due to H1 loss on the maternal genome. However, H1 loss differentially influences DNA methylation in heterochromatic and euchromatic TEs prior to DME action (Zemach et al., 2013), making it difficult to delineate if H1 directly or indirectly involves in DME demethylation in the central cell. Furthermore, DME and its more ubiquitous paralog ROS1 are likely recruited by and make contact with specialized local chromatin environments (Qian et al., 2012), as demonstrated by a recent study showing that ROS1 binds to all 4 histone proteins *in vitro* (Parrilla-Doblas et al., 2022). Thus, the interaction between H1 and DME might have more of a structural or physical rather than a regulatory purpose.

### H1.2 and DME are co-expressed in the central cell

In *Arabidopsis*, the reproductive phase is initiated late in adult plant with the specification of meiocyte precursors known as the spore mother cells (SMCs) that transition from vegetative to reproductive cell fate. This transition is accompanied by a temporary eviction of H1.1 and H1.2 in the megaspore mother cells, the female SMCs (She et al., 2013;She and Baroux, 2015;Ingouff et al., 2017). During male gametogenesis, both H1 variants are absent in late microspore stage, remain absent in the VC nuclei, but are present in the sperm nuclei (Hsieh et al., 2016;He et al., 2019). The absence of H1 in the VCs where DME is active indicates that H1 is not a requirement for DME demethylation, at least not in the VCs. By contrast, we found that the *H1*.*2* promoter is active in the central cell where DME is also active (**Figure 2**), highlighting the difference in chromatin organization among these two gamete companion cells where DME primarily functions. For instance, the FACT complex is specifically required for DME demethylation in the central cell heterochromatin, but not in the VC (Frost et al., 2018), suggesting that CC has a more compact nuclear structure than that in VC and supporting our finding that H1 variants are present and function to maintain epigenetics homeostasis in the central cell.

### H1 mutations did not cause a *dme*-like seed abortion phenotype

If H1 is required for normal DME function in the central cell, loss of H1 would be expected to induce certain degree of seed abortion similar to what’s seen in the *dme* mutant. We examined seed phenotype of *h1* single, double, and triple mutants (*h1*.*1-1 h1*.*2-2 h1*.*3-1*) using different batches of plants. Seed abortion rates of *h1* single mutants *h1*.*1-1, h1*.*2-2, and h1*.*3-1* were 1.80%, 1.61%, and 1.26%, respectively, which were slightly higher than that of wild type Col-0 (0.62%) (**Table S5**). The *h1* double mutants *h1*.*1-1 h1*.*2-2* and *h1*.*2-2 h1*.*3-1* clearly have a cumulative effect on seed abortion, which reach 6.71% and 2.80% respectively. The *h1* triple mutant (*h1*.*1-1 h1*.*2-2 h1*.*3-1*) has the highest seed abortion rate (7.35%) (**Table S5**), which shows statistically significant difference from any other *h1* single and double mutants except *h1*.*1-1 h1*.*2-2*. This result suggests that both *H1*.*1* and *H1*.*2* play a role in seed development. Interestingly, we also observed a big variation of seed abortion rate in histone *h1* mutants. In the *h1* triple mutant, the spectrum of seed abortion rate in individual silique varied from 0% to 75%, and more than 61% (113 out of 184) examined siliques have aborted seeds (**Table S6**). This big variation of seed abortion can reflect subtle epigenetic influence of H1 on seed development as it has now been shown that H1 affects flowering time, stomata and lateral root formation, and stress response (Rutowicz et al., 2019). This can also suggest that epigenetic effect of H1 on seed development might be easily influenced by environment or subtle plant growth conditions. In short, H1 loss has a subtle influence on seed development, but *h1* mutants do not exhibit a *dme*-like prominent seed abortion phenotype, and H1 is not a strict requirement for DME function in the central cell.

### Maternal *h1* triple mutant affects chromatin architects and influences DME demethylation

H1 binds to the nucleosome core and the linker DNA between two adjacent nucleosomes and is required for higher-order chromatin organization; yet, despite its ubiquitous presence in the nucleus and important nuclear functions, loss of H1 has very limited effects on gene transcription and TE silencing (Rutowicz et al., 2019). H1-containing nucleosomes are natural barriers to the enzymes that modify DNA, such as DNA methyltransferases (Lyons and Zilberman, 2017) (Zemach et al., 2013), DNA repair enzymes, and DNA demethylases (He et al., 2019) (Frost et al., 2018). Consequently, loss of H1 facilitates access of DNA methyltransferases to the decondensed heterochromatin and resulted in a gain in heterochromatic TEs methylation (Zemach et al., 2013). By contrast, euchromatin TEs exhibited a loss in DNA methylation in *h1* homozygous mutant, for reasons currently not fully understood (Zemach et al., 2013).

Our *h1* x Ler endosperm methylome data supports this general model of H1 function. Our results suggest that loss of H1 in the central cell allowed easier access of DME, particularly to the heterochromatic regions. This is reflected in a 2x increased in the number of EMB-ENDO DMRs in *h1*-wt hybrid compared with wild type seeds (**Figures 4A, 4B**), and the gained (*h1*-unique) DMRs are enriched for heterochromatin states (**Figure 4D**) and in the pericentromere regions (**Figure 4E**). Since there was little change between *h1*-wt and wild-type embryo methylome, we can assume the difference in endosperm methylome was caused by DME action in the *h1* central cell. Our data also suggests that H1-containing nucleosomes impede DME demethylation process and the removal of H1 resulted in a significant increase in DMR length (**Figure 4C**). This notion was supported by our earlier studies that the FACT complex is specifically needed in the H1-bound, nucleosome rich heterochromatin targets (Frost et al., 2018;Zhang et al., 2019). Within the canonical DME target loci (*dme* vs wild-type endosperm), we did not observe any significant difference between embryo and endosperm (**Figure 3A**, blue and green traces); but a small degree of difference is visible outside DME canonical loci with a slight hypomethylation in *h1*-wt endosperm (**Figure 3A**), consisting with gaining more DMRs in *h1* hybrid due to a more extensive demethylation in the *h1* CC genome. A direct comparison between *h1*-wt and wild-type endosperm revealed that loss of H1 differentially affect DME canonical target loci (**Figure 3C**, red trace). Canonical loci hypermethylated in *h1* endosperm (**Figure 3C**, positive side of the red trace) are enriched for genic coding sequences whereas loci hypomethylated (red trace, negative side) are more enriched for intergenic and heterochromatin sequences (**Figure 3D**). Taken together, our methylome data showed the effect of H1 loss on DME demethylation is largely indirect.

### H1 loss in the maternal genome has a very limited influence on gene transcription and imprinting regulation

Our attempt to isolate RNAs from FACS-purified endosperm nuclei was unsuccessful upon multiple trials with different RNA extraction protocols. This prevented us from performing a more detailed analysis on how loss of H1 influences gene imprinting regulation. Due to the presence of maternal seedcoat tissues, revealed by tissue enrichment test (**Figure S10**), we instead limited our allele-specific expression analysis on a list of previously reported high-confident imprinted genes (Schon and Nodine, 2017) and qualitatively assessed whether maternal H1 loss induced a significant disruption on their allele-specific expression. Very few differentially expressed genes or TEs were identified between *h1*-wt and wild type in endosperm and in embryo, suggesting that the wild-type H1 alleles from pollen can quickly compensate for the loss of maternal copies upon fertilization. We used the ratio of maternal over total transcripts as a measurement of parental bias and did not identify any deviation between *h1*-wt and wild type among known imprinted genes. Overall, the maternal-bias score for each imprinted gene were highly correlated between *h1* hybrid and wild type, with selected individual gene showing some variation visible on the scatter plot (**Figure 5B**). These observations showed that even with a greater degree of maternal genome demethylation in *h1*, the influences on endosperm gene transcription or imprinting regulation was very limited. However, we cannot rule out the possibility that H1 might play a more direct role in assisting DME demethylation at specific loci via post-translational modification.

We reported earlier the expression of *MEA, FIS2*, and *FWA* were reduced and their maternal copies were hypermethylated in *h1*-wt hybrid endosperm (Rea et al., 2012). Here we showed that the flanking sequences of these three genes are indeed hypermethylated in the *h1*-wt endosperm relative to wild type and except for *MEA* whose expressing was below detection in our RNA-seq datasets, the expression of *FIS2* and *FWA* was reduced in the *h1*-wt hybrid endosperm. This was independently validated by the promote::GFP reporter constructs showing *MEA* and *FWA* expression were reduced in *h1* endosperm compared with wild type. We believe this is due to the differential effect on DME target sequences in the absence of H1 (**Figure 3C**). However, we also observed a variable methylation change in other selected imprinted genes (**Figure S6**). The reason why maternal H1 loss caused hypermethylation in some genes but not others remains to be investigated. A recent study revealed that without H1, siRNA biogenesis is quantitatively expanded to non-CG-methylated loci, which might contribute to this disparate methylation change (Choi et al., 2021).

In summary, our data revealed that loss of H1 in the central cell resulted in a less condensed heterochromatin configuration which indirectly facilitates access of DME to its nucleosome targets.

## Data availability statement

The NGS datasets reported in this study can be found in NCBI GEO under GSEXXXX (in process).

## Author contributions

RLF, WX and TFH conceived the project; QH, YHH, CZ, AB, ME, HY, CP performed the experiments; XQZ, QH, YHH, WX, TFH analyzed the data; RLF, WX and TFH wrote the article with contributions of all the authors. All authors contributed to the article and approved the submitted version.

## Acknowledgments

We thank the Arabidopsis Biological Resource Center at Ohio State University for providing Arabidopsis mutant seed stocks, Dr. Grant Kolar and Caroline Murphy at Research Microscopy and Histology Core at SLU for providing assistance with confocal microscopy, Joy Eslick at Flow Cytometry Research Core Facility at SLU for helping with nuclei sorting.

