## Supplemental Figures and Tables for "Loss of Linker Histone H1 in the Maternal Genome Influences DEMETER-Mediated Demethylation and Affects the Endosperm DNA Methylation Landscape"

### Supplementary Figures

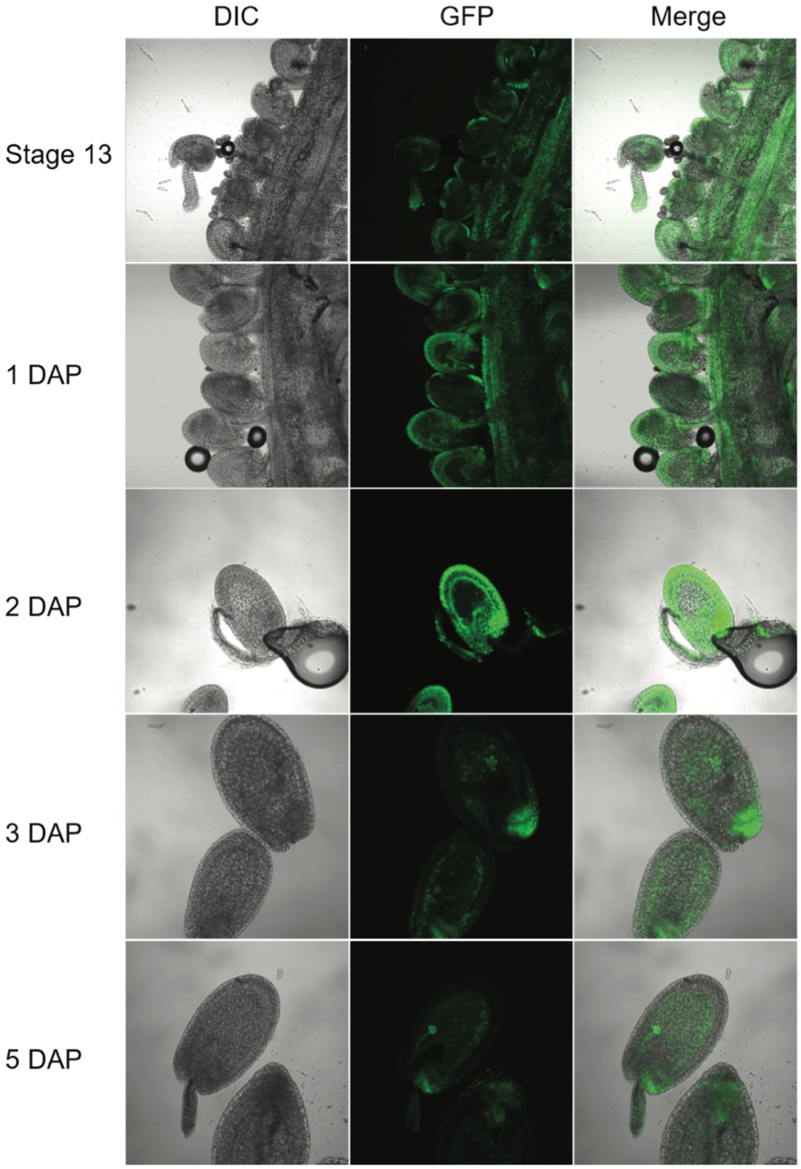


FIGURE S1. Expression of the *H1.1 promoter::GFP* transgene in *Arabidopsis* ovules and young developing seeds. Ovules and seeds with expression of *H1.1 promoter*:*GFP* transgene were photographed using confocal fluorescence microscope. Ovules at flower stage 12 and stage 13 and seeds at 1, 3, 5, and 7 DAP were hand-dissected for imaging. The GFP signal is shown in green. DAP, Days After Pollination; DIC, Differential Interference Contrast.

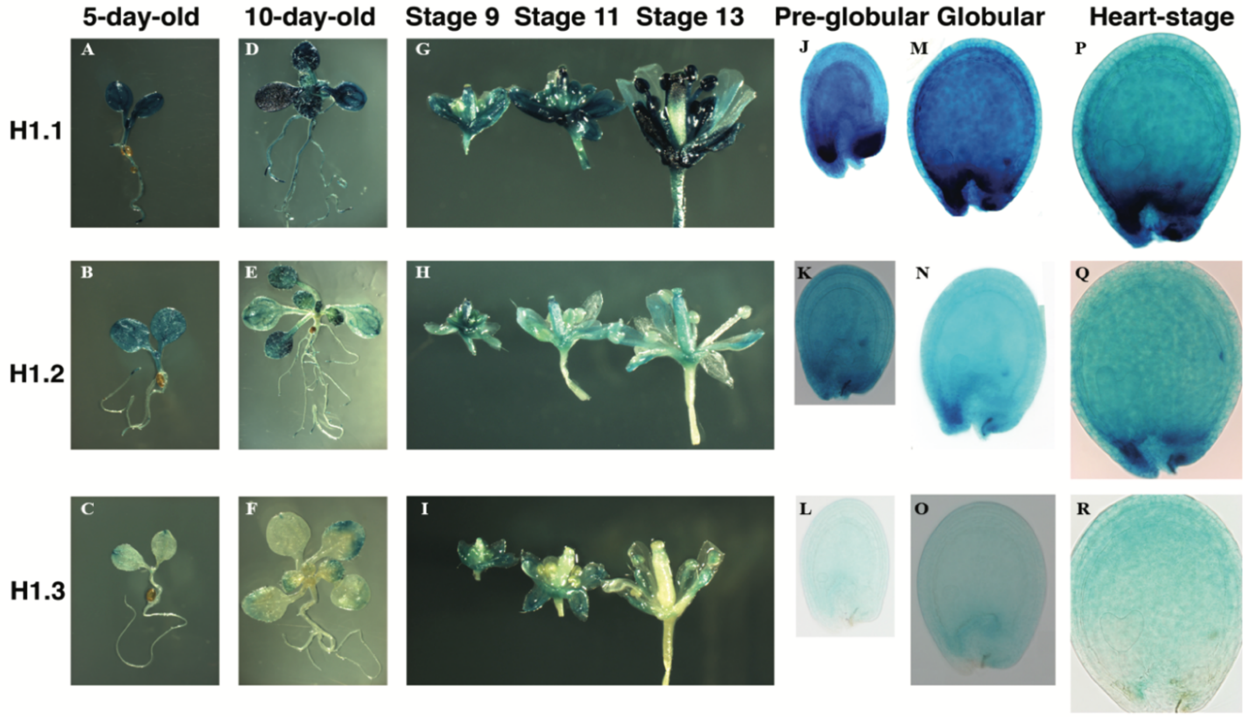


FIGURE S2. Expression of the three *H1 promoter::GUS* reporters in various *Arabidopsis* tissues. (A-C) 5-day-old young seedlings after 3-day cold treatment. (D-F) 10-day-old young seedlings after 3-day cold treatment. (G-I) Side views of opened whole flowers at stages 9, 11, and 13, respectively. (J-L) Seeds photographed when embryos are at Pre-globular stage. (M-O) Seeds photographed when embryos are at globular stage. (P-R) Seeds photographed when embryos are at Heart stage.

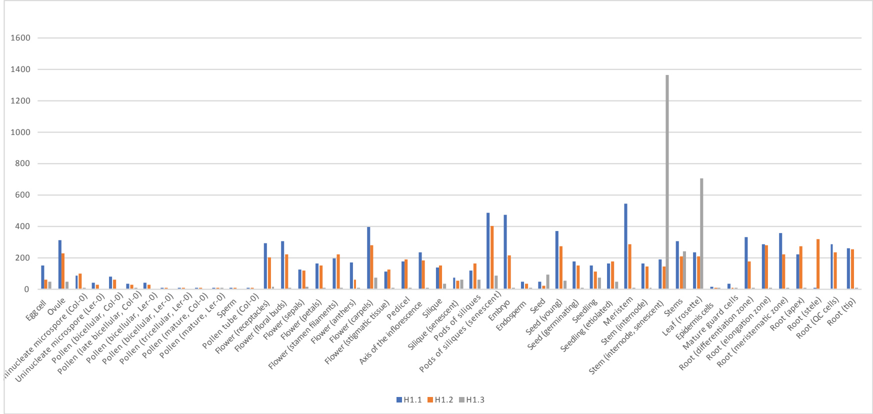


FIGURE S3. Expression patterns of H1.1, H1.2, and H1.3 across tissues and developmental stages**.** Raw expression values are TPM normalized retrieved from the Evorepro database.

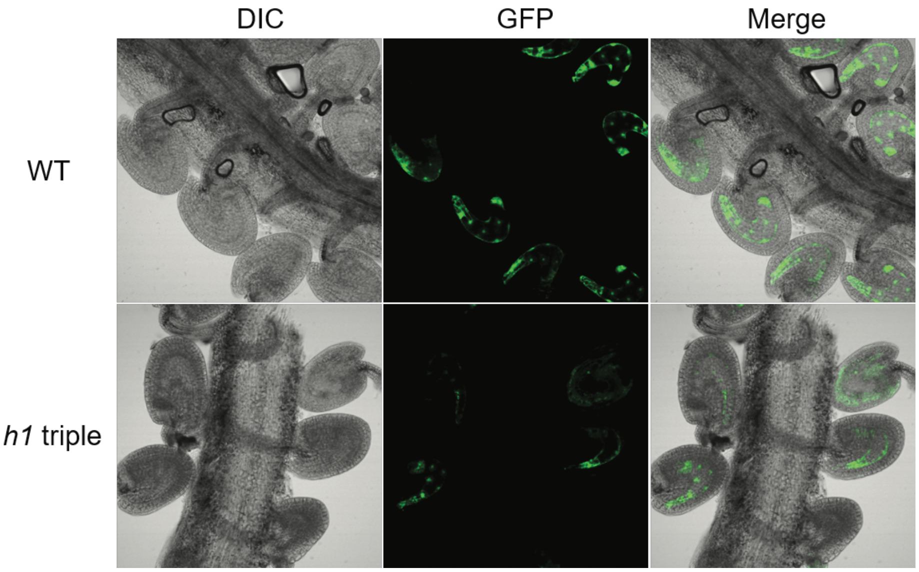


FIGURE S4. Expression of the transgene *pFWA:GFP* is reduced in the endosperm in the *h1* triple mutant. Ovules and seeds with *pFWA:GFP* transgene are photographed using confocal fluorescence microscope. Seeds at 1 DAP are dissected out for imaging. GFP expression is shown in green. DAP, Days After Pollination; DIC, Differential Interference Contrast.

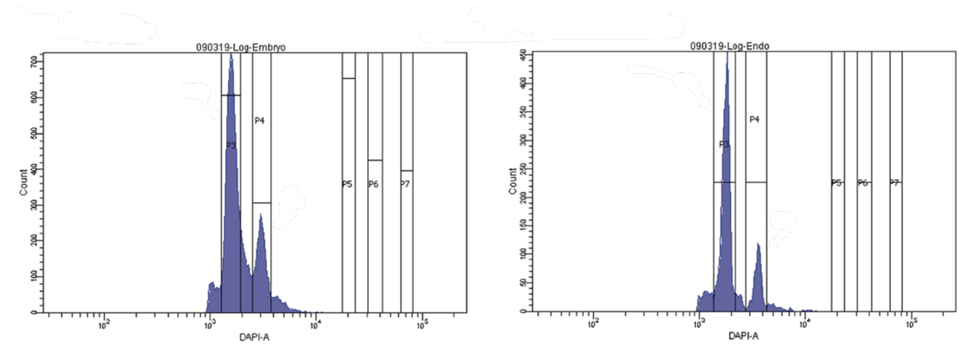


FIGURE S5. Flow cytometric profiles showing DNA contents of nuclei with different DAPI measurements. Nuclei were extracted from dissected embryo (A) and endosperm (B) derived from crosses between Col and L*er* at 7 days after pollination. The ploidy of collected peak is indicated.

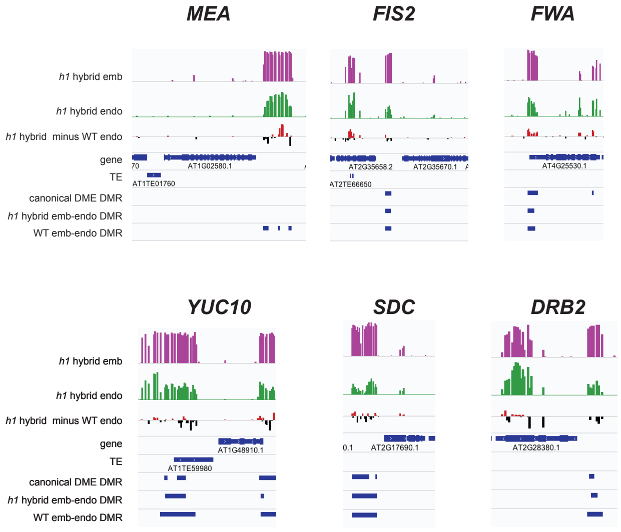


FIGURE S6. Selected imprinted genes hyper- or hypo-methylated in h1-wt hybrid endosperm. Genome browser snapshots of imprinted genes examples hyper- (top panel) or hypo- (bottom panel) methylated in *h1*-wt hybrid endosperm. CG methylation difference between *h1*-wt and wt endosperm is shown in the third track. The positions of three sets of DMRs are indicated as blue boxes in the bottom three tracks.

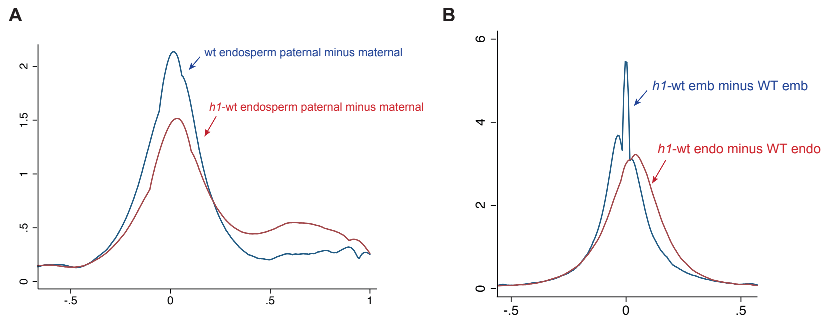


FIGURE S7. (A) Parental genome CG methylation difference in endosperm. Kernel density plot showing the differentiation in parental CG DNA methylation (paternal minus maternal) in wild type (blue) and *h1*-wt hybrid (red) endosperm. (B) Parental genome CG methylation difference in endosperm. Kernel density plot of CG methylation difference between wild-type and *h1*-wt hybrid embryo (blue) and endosperm (magenta).

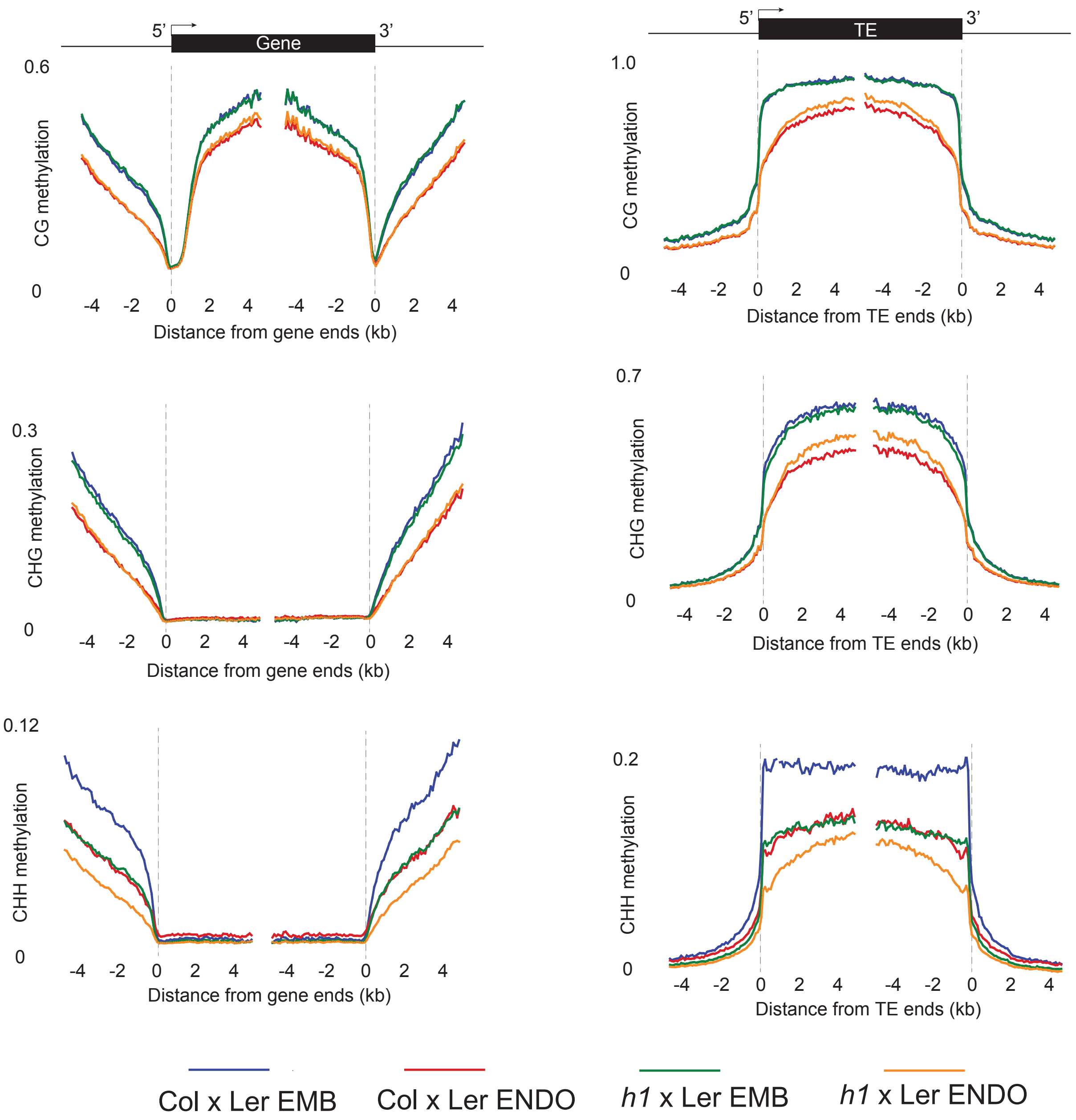


FIGURE S8. Whole genome CG, CHG, CHH methylation metaplots. Average CG, CHG, CHH methylation in Genes (left panels) or TEs (right panels) of embryo and endosperm isolated from wild type and *h1*-wt hybrid seeds.

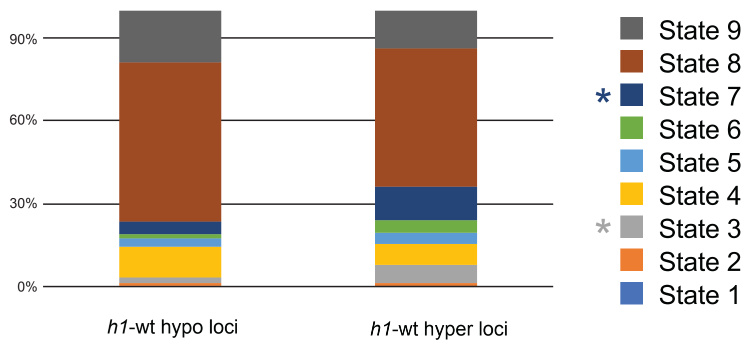


FIGURE S9. Different chromatin states in the canonical DME targets in endosperm. Percent distribution of 9 different chromatin states in the canonical DME targets that loose (*h1*-hypo) or gain (*h1*-hyper) in *h1*-wt hybrid endosperm compared with wild type.

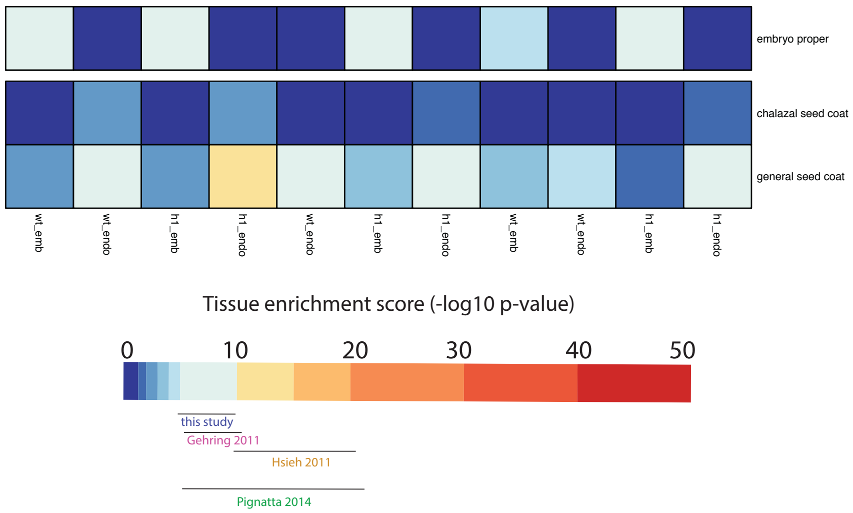


FIGURE S10. Assessing the degree of maternal RNA contamination by tissue enrichment test. Columns represent different RNA-seq transcriptomes used in this study, and rows represent two seed coat tissues. Our endosperm transcriptomes are moderately enriched for some seedcoat-specific transcripts.

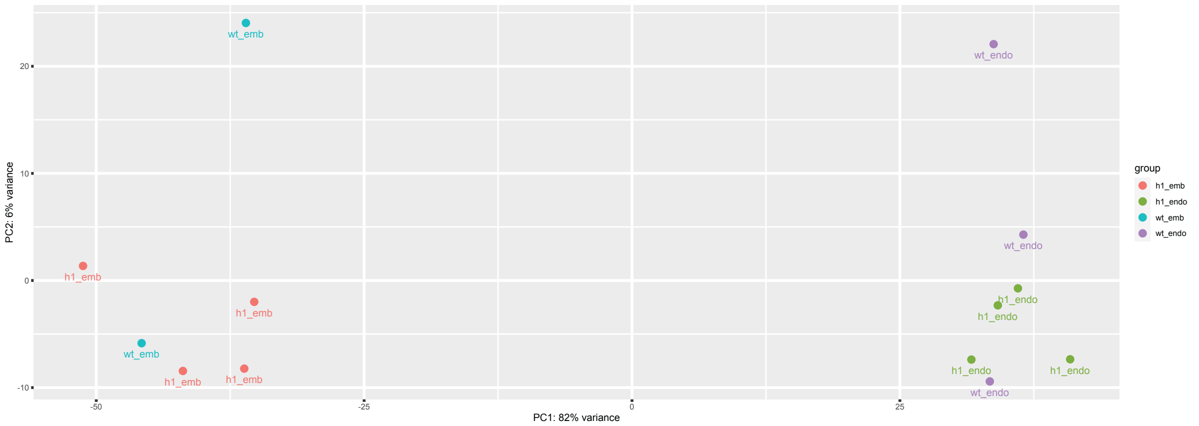


FIGURE S11. PCA analysis of all 13 samples used in this study.

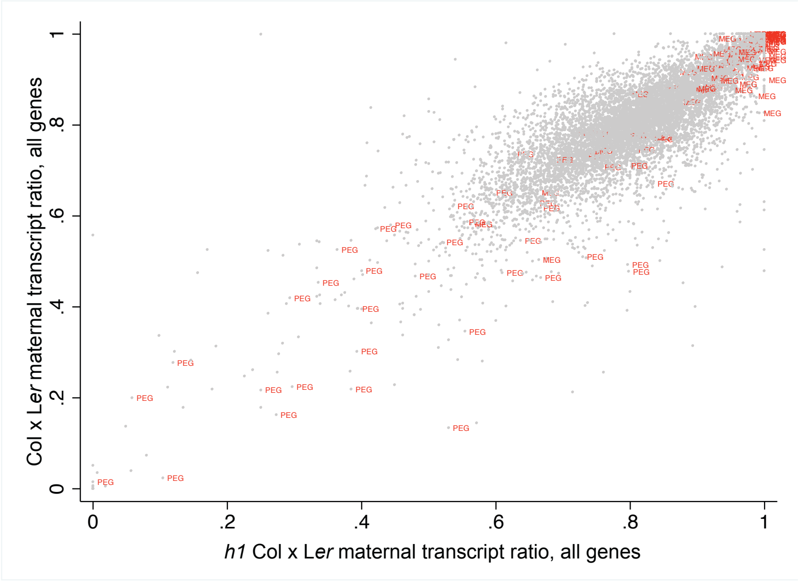


FIGURE S12. Correlation of maternal transcript proportion between wild type and *h1*-wt hybrid endosperm. Scatterplot showing the correlation of maternal transcript proportion of all genes between wild type and *h1*-wt hybrid endosperm.

### Supplementary Tables

#### Supplemental Table 1. BS-seq dataset information

| Sample | Unique mapped reads | Coverage | BS conversion rate* | M/P ratio |
|---|---|---|---|---|
| *h1*xLer emb rep1 | 12781829 | 7.7 | 99.76% | 1.055 |
| *h1*xLer endo rep1 | 22099703 | 13.3 | 99.74% | 2.275 |
| *h1*xLer emb rep2 | 35405734 | 21.2 | 99.78% | 1.04 |
| *h1*xLer emb rep3 | 15696292 | 9.4 | 99.66% | 1.07 |
| *h1*xLer endo rep2 | 39038514 | 23.4 | 99.78% | 2.31 |
| *h1*xLer endo rep3 | 28919583 | 17.4 | 98.82%** | 2.04 |

*Bisulfite conversion rate was calculated using chloroplast DNA methylation ratio.
**Excluded from further analysis due to low conversion rate.

#### Supplemental Table 2. Whole genome *h1* x Ler embryo and endosperm BS-seq data correlation between bio-replicates.

| Pearson correlation coefficient | *h1* emb rep1 $^{m}CG$ | *h1* emb rep2 $^{m}CG$ | *h1* emb rep3 $^{m}CG$ |
|---|---|---|---|
| *h1* emb rep1 | 1 | | |
| *h1* emb rep2 | 0.9516 | 1 | |
| *h1* emb rep3 | 0.9243 | 0.9487 | 1 |

| Pearson correlation coefficient | *h1* endo rep1 $^{m}CG$ | *h1* endo rep2 $^{m}CG$ | *h1* endo rep3 $^{m}CG$ |
|---|---|---|---|
| *h1* endo rep1 | 1 | | |
| *h1* endo rep2 | 0.9563 | 1 | |
| *h1* endo rep3 | 0.9157 | 0.9331 | 1 |

#### Supplemental Table 3. Pair-wise overlap between different sets of DMRs.

| | SP vs VC | *dme* vs wt EN | wt_EM_EN | *h1*_EM_EN |
|---|---|---|---|---|
| SP vs VC | 100% | | | |
| *dme* vs wt EN | 51% | 100% | | |
| wt_EM_EN | 34% | 56% | 100% | |
| *h1*_EM_EN | 80% | 84% | 86% | 100% |

**Supplemental Table 4. RNA-seq data correlation between bio-replicates.**

| Pearson correlation coefficient | wt_endo rep1 | wt_endo rep2 | wt_endo rep3 |
|---|---|---|---|
| wt_endo rep1 | 1 | | |
| wt_endo rep2 | 0.8355 | 1 | |
| wt_endo rep3 | 0.9489 | 0.8948 | 1 |

| Pearson correlation coefficient | wt_emb rep1 | wt_emb rep3 |
|---|---|---|
| wt_emb rep1 | 1 | |
| wt_emb rep3 | 0.6583 | 1 |

| Pearson correlation coefficient | *h1*_endo rep1 | *h1*_endo rep2 | *h1*_endo rep3 |
|---|---|---|---|
| *h1*_endo rep1 | 1 | | |
| *h1*_endo rep2 | 0.9302 | 1 | |
| *h1*_endo rep3 | 0.7789 | 0.9321 | 1 |

| Pearson correlation coefficient | *h1*_emb rep1 | *h1*_ emb rep2 | *h1*_ emb rep3 |
|---|---|---|---|
| *h1*_ emb rep1 | 1 | | |
| *h1*_ emb rep2 | 0.9778 | 1 | |
| *h1*_ emb rep3 | 0.9863 | 0.941 | 1 |

**Table S5**. Histone H1 mutation causes seed abortion.

| **Genotype** | **Total Seeds** | **% Abortion** | **Total Seeds Per Silique** |
|---|---|---|---|
| **Wild Type Col-0** | 3366 | 0.62% | 52.6 |
| **Wild Type L*er*** | 444 | 0.90% | 49.3 |
| ***h1.1-1*** | 3664 | 1.80% | 54.7 |
| ***h1.2-2*** | 1676 | 1.61% | 40.9 |
| ***h1.3-1*** | 2530 | 1.26% | 53.8 |
| ***h1.1-1 h1.2-2*** | 2534 | 6.71% | 46.9 |
| ***h1.1-1 h1.3-1*** | 4372 | 1.88% | 49.1 |
| ***h1.2-2 h1.3-1*** | 2145 | 2.80% | 45.6 |
| ***h1.1-1 h1.2-2 h1.3-1*** | 8362 | 7.35% | 45.4 |

Note: % Abortion is the percentage of aborted seeds in total examined seeds. % Good is the percentage of viable seeds in total examined seeds.

**Supplemental Table S6.** Distribution and variation of seed abortion among siliques in wild type and the *h1* triple mutant.

| Seed abortion rate / Genotype | 0% | 0.1-4.9% | 5-9.9% | 10-19.9% | 20-29.9% | 30-39.9% | >40% | total |
|---|---|---|---|---|---|---|---|---|
| **WT Col-0** | 48 | 14 | 2 | 0 | 0 | 0 | 0 | 64 |
| ***h1.1-1 h1.2-2 h1.3-1*** | 71 | 40 | 25 | 22 | 15 | 7 | 4 | 184 |

Note: the table shows the number of siliques with different seed abortion rates among total examined siliques. Seed abortion rate is the percentage of aborted seeds in each silique.
